# DeepVASP-E: A Flexible Analysis of Electrostatic Isopotentials for Finding and Explaining Mechanisms that Control Binding Specificity

**DOI:** 10.1101/2021.08.22.456843

**Authors:** Felix M. Quintana, Zhaoming Kong, Lifang He, Brian Y. Chen

## Abstract

Amino acids that play a role in binding specificity can be identified with many methods, but few techniques identify the biochemical mechanisms by which they act. To address a part of this problem, we present DeepVASP-E, an algorithm that can suggest electrostatic mechanisms that influence specificity. DeepVASP-E uses convolutional neural networks to classify an electrostatic representation of ligand binding sites into specificity categories. It also uses class activation mapping to identify regions of electrostatic potential that are salient for classification. We hypothesize that electrostatic regions that are salient for classification are also likely to play a biochemical role in achieving specificity. Our findings, on two families of proteins with electrostatic influences on specificity, demonstrate that large salient regions can identify amino acids that have an electrostatic role in binding, and that DeepVASP-E is an effective classifier of ligand binding sites.

## 1. Introduction

A small minority of amino acids play central roles in selective binding. Discovering those amino acids, and especially the biochemical mechanisms by which they act, is crucial for understanding how genetic variations influence pathogenicity and for interpreting how preferred binding partners might be changed through protein redesign. Most approaches proposed to date have focused on identifying influential amino acids. Evolutionary techniques^1,2^ infer that the conservation of amino acids, or variations that follow major evolutionary divergences, are evidence for a role in function. Cavity based techniques^3–5^ infer that proximity to the largest clefts on the solvent accessible surface is evidence for an enriched role in function. Structure comparison algorithms^6–8^ infer that having certain atoms or amino acids in specific geometric configurations is evidence for the capacity to catalyze the same chemical reaction. Combinations of these and other concepts have also been considered.^9,10^ All these methods can focus human attention on amino acids that may have a functional role, thereby reducing effort wasted on irrelevant amino acids. However, without deducing the biochemical mechanism by which these amino acids contribute to selective binding, the problem of determining the biochemical effect of genetic variation, and the design of validation experiments, must still rely on human expertise. Given that mutations at only a few critical residues have combinatorial effects on function, a computer generated explanation of specificity mechanisms could offer insights at appropriate scale and suggest mechanisms that experts might overlook.

Towards interpreting the biochemical role of individual residues, this paper proposes a novel general strategy that we call the *Analytic Ensemble* approach, and it examines one aspect of this strategy. We begin with a *training family* of closely related proteins that perform the same biochemical function, with subfamilies that prefer to act on similar but non-identical ligands. These families could be evolutionary in origin, or simply groups of mutants that share binding preferences. Suppose that we design an algorithm that examines only patterns of steric hindrance to identify amino acids that influence specificity. Then any amino acid identified by this narrow approach can be associated with a steric influence on specificity, because the method examines no other mechanism. Imagine a second approach that examines only electrostatic fields to find influential amino acids. Residues found by the second approach must have an electrostatic influence on specificity for the same reasons. Further methods could be developed for hydrogen bonds, hydrophobicity, and so on. While these individual approaches are very narrow, they could collectively analyze a diverse range of structural mechanisms, and their exclusivity has the novel property that it connects their findings to a biochemical mechanism. The same inferential structure is typically not possible with existing approaches because most employ biochemically holistic representations. For example, finding the same residues in the same locations with a structure comparison algorithm could identify amino acids that might be ideally positioned to either form hydrogen bonds, or to sterically hinder a discouraged ligand. In such cases, important residues can be detected, but activity through steric hindrance, for example, cannot be confirmed.

As a part of the Analytic Ensemble, this paper presents DeepVASP-E, an algorithm for identifying electrostatic mechanisms by which amino acids influence specificity. It achieves this purpose by representing the electric field within protein binding sites using a voxel representation of electrostatic isopotentials. Given isopotentials *q* derived from the binding site of a query protein with unknown binding preferences, DeepVASP-E performs two functions: First, it adapts a three dimensional convolutional neural network (3D-CNN) to classify *q* into one of the subfamilies based exclusively on the geometric similarity of electrostatic isopotentials. Second, it uses the gradient-weighted class activation mapping, Grad-CAM++^11^ to identify regions of *q* that motivate its classification into that subfamily. We hypothesize that regions identified in this way will identify zones of electrostatic potential that are important for selective binding, thereby proposing a simple electrostatic mechanism by which the query protein achieves specificity.

Deep learning methods have recently made increasing contributions to structural bioinformatics, most notably for the prediction of protein structures.^12–15^ DeepVASP-E has some similarities to these methods in its underlying techniques. For example, 3D-CNNs have been applied for the prediction of binding sites^16,17^ and the classification of proteins by Enzyme Classification number.^18^ Grad-CAM maps have also been applied for the identification of functional amino acids.^19^ While it also employs deep learning techniques, DeepVASP-E is using deep learning for a new purpose, to generating biochemical explanations for specificity.

Our attention to electrostatic fields in DeepVASP-E is inspired by findings with VASP-E, a volumetric algorithm for comparing and analyzing electrostatic isopotentials in ligand binding sites and protein-protein interfaces.^20,21^ VASP-E uses Constructive Solid Geometry (CSG), a technique for computing unions, intersections and differences of geometric solids, to perform these comparisons. Fig. 1 shows an example of the electrostatic properties of a binding cavity.

**Fig. 1.**
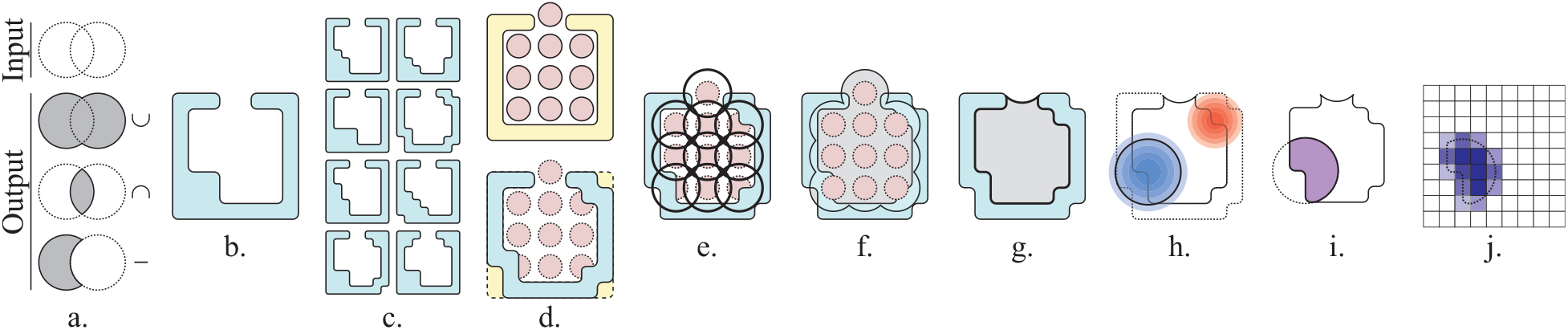
Representing the Electrostatic Properties of a Binding Cavity. **a**. CSG union (∪), intersection (∩), and difference operations (−) performed on input solids (white) and their outputs (grey). **b**. A protein illustrated as a molecular surface (teal). **c**. Conformational samples of the protein. **d**. The pivot structure *t*′ (top, yellow) with ligand atoms (red), and the structural alignment (bottom) of one conformational sample (teal) to *t*′. **e**. Spheres defining the neighbourhood of the ligand atoms (black circles). **f**. The union of the ligand atom spheres (black outline) minus the molecular surface of the conformational sample (teal), is shown in grey. **g**. The binding site in the conformational sample (grey). **h**. The positive (blue gradient) and negative (red gradient) regions of the electrostatic field, shown with the selected electrostatic isopotential (black circle). **i**. The cavity field: the CSG intersection between the electrostatic isopotential and the cavity region (purple). **j**. Translation of the cavity field (dotted outline) into the weighted voxel representation for 3D-CNN training (shaded purple squares).

We found that comparisons of electrostatic isopotentials within binding sites, represented as geometric solids, could classify proteins that prefer different ligands and predict amino acids that influence specificity.^22^ This paper builds on this technique by introducing a deep learning approach that can be interpreted to identify the regions of the electrostatic field that are salient to classification.

To perform classification and saliency analysis accurately, it is vital to compensate for conformational variations that may be observed in the training family. While proteins can occupy a wide range of conformations, common sidechain motions and backbone breathing can alter the apparent solvent accessibility of a binding cavity and the relative positions of its active residues. For this reason, DeepVASP-E uses conformational samples of the training family from medium-timescale molecular dynamics simulations to train the neural network. Our integration with simulated data has two novel and synergistic advantages: First, it provides a source of highly authentic conformational samples for training that can assist in compensating for conformational change. Second, regarding the intensive need for training data in deep learning systems, simulations offer a prolific source of augmented data that can mitigate the scarcity of experimentally determined structures.

Our results examine the performance of DeepVASP-E on two sequentially nonredundant families of proteins with experimentally established binding preferences. We evaluated how accurately the method classifies binding cavities based on electrostatic isopotential data alone, and compared its classification performance to existing techniques. We then validated the accuracy of the saliency maps against experimentally established specificity mechanisms. Together, these results point to novel applications in deconstructing protein function from structure.

## 2. Methods

### Overview

DeepVASP-E uses a training family *T* constructed from a family of proteins with well defined subfamilies *{T*_0_, *T*_1_, …*T*_*n*_*}* that exhibit distinct binding preferences and a known ligand binding site. To prepare this data, as summarized in Fig. 1, the structure of each protein in *T* is first simulated using molecular dynamics to produce conformational samples (Fig. 1c). Second, each conformational sample, *t*, is structurally aligned to a *pivot structure t*′, which is a member of *T* that was chosen because it has a ligand crystallized in the binding site (Fig. 1d). We use the ligand atoms, now aligned to the binding site of *t* to define the binding site (Fig. 1e-g). Third, we determine the electrostatic field, and given an isopotential threshold, we compute an electrostatic isopotential of *t* (Fig. 1h). Using CSG, regions of the isopotential that are outside the binding site are removed, producing a region we call the *cavity field* (Fig. 1i). Finally, the cavity field is translated into voxel data, as input for the 3D-CNN (Fig. 1j).

When classifying the structure of a novel query protein *q* into one of the *n* subfamilies, we treat it as a single conformation of the novel protein. Thus, to prepare *q* for classification, we begin with the structural alignment of *q* to *t*′ and follow the data preparation steps above until voxel data is generated. Using a 3D-CNN model that has been trained, we produce classifications of *q* into one of the *n* subfamilies (Fig. 2).

**Fig. 2.**
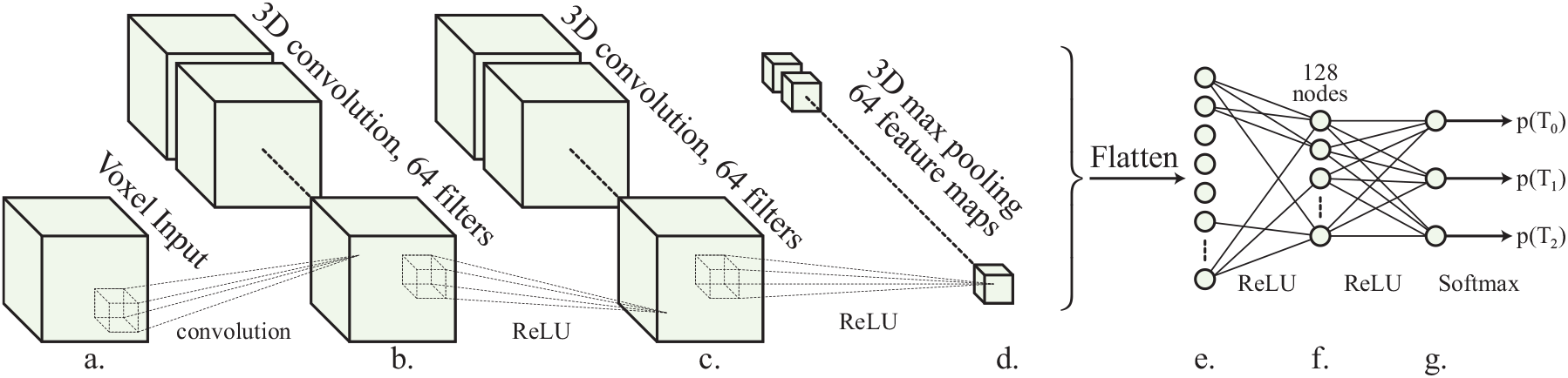
**a**. Real valued voxel inputs with each of the three dimensions ranging from 30 - 42 cubes. **b, c**. 3D convolutional layers with 64 filters. 5×5×5 kernels with stride 1 were used (dotted lines), with padding to maintain resolution. A ReLU activation function is used in both layers. **d**. 3D max pooling layer with pool size 2×2×2 and stride (2,2,2), producing outputs of size (*x/*2, *y/*2, *z/*2) with ReLU activation function. **e**. Flattening layer with ReLU activation function. **f** Fully connected layer reducing layer e to 128 notes, ReLU activation layer. **g**. output layer with 3 nodes corresponding to the number of classes with softmax activation function, to produce probabilities of classification

To predict the voxels that are electrostatically significant for selective binding, we adapt the gradient-weighted class activation map method Grad-CAM++ to identify the regions most associated with classification.^11^ The resulting voxels define binding cavity regions that are significant for classification. We hypothesize that these differentiating potentials are nearby amino acids that are crucial for binding, which we will verify against experimentally established results in Section 3.

### Training Families

To test DeepVASP-E, we selected the serine protease and enolase superfamilies as training families (Table 1). Within each training family, we selected three distinct subfamilies with different binding preferences. From the serine proteases, we selected the trypsin, chymotrypsin, and elastase subfamilies, and we selected the enolase, mandelate racemase, and muconate lactonizing enzyme families from the enolase superfamily.

**Table 1.** PDB codes of training families used in this study.

| Serine Protease Superfamily |  |  | Enolase Superfamily |  |
| --- | --- | --- | --- | --- |
| Trypsins | 1a0j, 1ane, 1aq7, 1bzx, 1fn8, 1h4w, 1trn, 2eek, 2f91 |  | Enolases | 1ebh, 1iyx, 1te6, 3otr |
| Chymotrypsins | 1ex3 |  | Mandelate Racemases | 1mdr, 2ox4 |
| Elastases | 1b0e, 1elt |  | Muconate Lactonizing Enzyme | 2pgw |

The serine proteases selectively cleave peptide bonds by recognizing amino acids on both sides of the scissile bond. The P1 residue, which is immediately before the bond, is recognized by the S1 specificity pocket. In Chymotrypsins, P1 is preferred to be large and hydrophobic.^23^ Trypsins prefer P1 residues that are positively charged, to complement their negatively charged S1 pocket.^24^ Elastases recognize a small hydrophobic P1 residues.^25^

The enolase superfamily share a binding site at the center of a TIM-barrel fold with an N-terminal “capping domain”. Members of this superfamily achieve a range of different functions that generally abstract a proton from a carbon adjacent to a carboxylic acid:^26,27^ Enolases catalyze the dehydration of 2-phospho-D-glycerate to phosphoenolpyruvate,^28^ mandelate racemases convert (R)-mandelate to and from (S)-mandelate,^29^ and muconate lactonizing enzymes catalyze the reciprocal cycloisomerization of cis,cis-muconate and muconolactone.^26^

The structures in our training families were selected from the Protein Data Bank^30^ (PDB) on 6.21.2011. Using the Enzyme Commission classifications (EC number), we found 676 serine protease and 66 enolase structures among the families selected for our data set. From this group, proteins with mutations, disordered regions, or enolases in closed or partially closed (and thus inactive) conformations were removed. From the remaining set, a set of sequentially nonredundant representatives were selected such that no representative had greater than 90% sequence identity to any other representative. Technical problems with molecular dynamics simulation prevented 8gch, 1aks, and 2zad from being included in this set. From each of the remaining structures we removed waters, ions, hydrogens, and other non-protein atoms.

### Conformational Sampling

To produce conformational samples of each protein, we used GROMACS 4.5.4.^31^ In preparation, each structure was centered in a cubic waterbox with 10 Å to the nearest point on the box. Inside, solvent was populated using an equilibrated 3-site SPC/E solvent model.^32^ Charge balanced sodium and potassium ions were added at a low concentrate (*<* 0.1% salinity).

Next, we performed Isothermal-Isobaric (NPT) equilibration of this system in four 250 picosecond timesteps, using a steepest descent algorithm. Starting with a position restraint force of 1000 kJ/(mol*nm), each step reduced the restraint by 250 kJ/(mol*nm). System energies were generated at the start of this equilibration, with initial temperature set at 300K and initial pressure at 1 bar. The Nosé-Hoover thermostat^33^ was used for temperature coupling. P-LINCS^34^ was used to update bonds. Electrostatic interaction energies were calculated by particle mesh Ewald summation (PME).^31^ The Parrinello-Rahman algorithm was used for pressure coupling.^35^ Temperature and pressure scaling were performed isotropically. The atomic positions and velocities of the final equilibration step were used to start the primary simulation, with all position restraints removed.

The primary simulation was sustained for 100 nanoseconds in 1 femtosecond timesteps. P-LINCS and PME were chosen for their parallel efficiency, and OpenMPI was used for inter-process and network communication. Simulations were run on multiple nodes with 16 cores each, with PME distribution automatically selected by GROMACS. The trajectory file of each completed simulation was then converted into individual timesteps in the PDB file format, with waterbox atoms removed. Each timestep was then rigidly superposed onto the original structure by minimizing RMSD between identical atoms [55]. From these timesteps, we selected 600 conformational samples at uniform intervals, and used them to train DeepVASP-E.

### Structural Alignment and Binding Site Representation

After we generated conformational samples of every protein in a training set, each one was aligned to the pivot structure using ska.^36^ Among the serine proteases, the pivot was bovine chymotrypsin (pdb: 8gch), and for enolases, the pivot was pseudomonas putida mandelate racemase (pdb: 1mdr). Pivot proteins were selected because they are co-crystallized with a ligand, which is used to localize the binding site in all conformational samples (Fig. 1e). Due to technical errors in MD simulation, 8gch was used only for this localization step.

After alignment we apply a technique from VASP, described earlier,^37^ to produce a solid representation of the binding site in the conformational sample. Paraphrasing here, we begin by producing spheres centered on the atoms of the ligand, with radius 5.0 Å (Fig. 1e). The CSG union of the spheres (Fig. 1f) defines the neighborhood of the ligand, *U*. We also compute a molecular surface *S* and envelope surface *E* of the conformational sample (not shown in Fig. 1f for clarity) using the classic rolling probe technique.^38^ Here, *S* and *E* are produced with probes with radius 1.4 Å and 5.0 Å respectively. *E* represents the region inside the protein, including the cavity. Using CSG, we compute (*U − S*) *∩ E* to produce the cavity (Fig. 1g).

### Computing Cavity Fields

Beginning with each conformational sample, we first model the position of all hydrogen atoms using the “reduce” component of MolProbity.^39^ We then use DelPhi^40^ to solve the Poisson-Boltzmann equation, producing the electrostatic potential field nearby (Fig. 1h). Finally, using isopotential thresholds -1.0 kt/e and 1.0 kt/e, we use VASP-E to compute electrostatic isopotentials from this field, representing each as a geometric solid. These thresholds were selected because in past experiments we considered a range of thresholds these values produced the clearest outputs.^20–22^ Higher absolute values create smaller isopotentials with less detail, while lower absolute values can be too large. Finally, we compute the CSG intersection between each isopotential and the cavity region defined above to produce a positive and a negative cavity field. Only positive cavity field is shown in Fig. 1i for clarity. Henceforth we perform separate computations on positive and negative cavity fields.

### Voxelized Binding Site Representations

Each cavity field is then translated into a voxel representation for 3D-CNN classification (Fig. 1j). First, we produce a bounding box around all positive or all negative cavity fields from all proteins in the training family. We then divide the bounding box into voxel cubes that are 0.5 °A on a side, padding it slightly to ensure an integer number of cubes in all dimensions. Finally, we use CSG intersections to estimate the volume in cubic angstroms of the cavity field inside each voxel. The vector of voxel intersection volumes is then passed into the 3D-CNN for training or classification.

### 2.1. Convolutional Neural Network

Our 3D-CNN architecture (Fig. 2) accepts voxel data as input. The architecture of DeepVASP-E is inspired by LeNet-5, a classic CNN architecture for recognizing handwritten and printed characters,^41^ and VoxNet, a 3D-CNN method for recognizing 3D point clouds.^42^ The chief design constraint for the CNN component of DeepVASP-E is the three dimensional resolution of the cavity field, which ranges between 30 to 42 cubes in each dimension, leading to a large number of neurons per layer that must be trained relative to typical 2D image analysis methods. Unfortunately, we have observed that reducing the number of neurons by using a coarser resolution can interfere with classification accuracy.^37^ For this reason we support the full resolution of the input cavity fields and a shallower topology similar to these classic methods. The approach concludes with a fully connected layer with softmax activation function to three categories corresponding to the three subfamilies of our training familes.

#### Class Specific Saliency Mapping

Gradient-weighted^++^ class activation mapping (GradCAM++)^11^ was used to generate saliency maps for all of the proteins and respective classes, which is a popular method for explaining CNN predictions. It uses the gradient information flowing into the last convolutional layer of the CNN to extract the importance of each neuron for the model output. Given a voxel’s spatial location (*x, y, z*) for a particular class *c*, we use Grad-CAM++ to generate a class-specific saliency map *L*^*c*^ as:

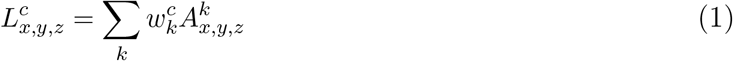

where 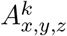 is the *k*th feature map in the last convolutional layer of 3D-CNN, and 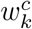 is the corresponding weight defined as follows:

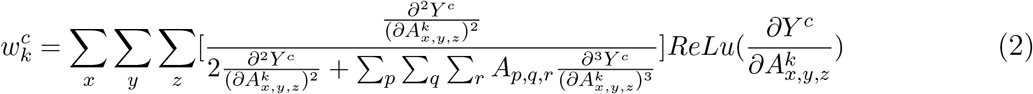

where *Y* ^*c*^ = exp(*S*^*c*^) is the class score, and *S*^*c*^ is the penultimate layer score of class *c* (Fig. 2g). 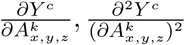 and 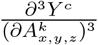 are the first-, second-, and third-order gradients w.r.t. 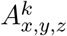

### 2.2. Experimental Design

To train the 3D-CNN model, each protein in the training family contributes 600 conformational samples to the data set. First, we leave all samples of one protein, the *evaluation set*, out of the dataset. Second, from the remaining samples, we randomly select 10% to create a *test set* to measure model classification accuracy. Next, the remaining snapshots are divided randomly into a *validation* set and *training* set at a 1:4 ratio. Weights on each node of the 3D-CNN are assigned and validated with the training and validation sets. This process repeats in each epoch until accuracy on the validation set stabilizes. We perform this process 10 times with a new and randomly selected test set. Finally, the weights of the model with the highest accuracy of all ten folds are used to predict the subfamily category of the samples in the evaluation set, which contains conformational samples from a protein that is not part of any other set. This evaluation is repeated, iteratively leaving out each protein in the training family.

### 2.3. Comparison with Existing Methods

While no other methods currently predict biochemical mechanisms that affect specificity, we compared the classification accuracy of DeepVASP-E to that of classic principal component analysis (PCA).^43^ For PCA, we learn a low-dimensional feature embedding of input data, and the embedding dimension is selected from {5, 10, 15, …, 100} using the same cross-validation strategy described in Section 2.2, then logistic regression model is applied as classifier.

#### Implementation Details and Availability

The deep learning backend is tensorflow-gpu (2.4.1) with Python 3.8. All experiments was performed on a workstation with 16 cores and 32GB main memory, using an Nvidia RTX 3090 GPU with 24GB of VRAM memory. Training associated with the 10-fold validation of a single protein in the dataset required approximate maximum of one hour with this machine.

## 3. Results

### Specificity Classification

We evaluated the performance of DeepVASP-E for predicting the specificity category of each member of both training families. The average Accuracy and F1-socre of compared methods are presented in Table 2.

**Table 2.** Comparison of classification results (avg *±* std).

| Training Family | Metric | PCA | DeepVASP-E |
| --- | --- | --- | --- |
| Serine Proteases<br>Positive Isopotential | Accuracy | 62.33 ± 26.73 | **96.95 ± 4.36** |
| | F1 | 72.58 ± 24.95 | **98.38 ± 2.32** |
| Serine Proteases<br>Negative Isopotential | Accuracy | 68.10 ± 40.18 | **100.00 ± 0.00** |
| | F1 | 71.05 ± 41.46 | **100.00 ± 0.00** |
| Enolase Superfamily<br>Positive Isopotential | Accuracy | 37.58 ± 14.17 | **97.02 ± 5.30** |
| | F1 | 52.53 ± 16.65 | **98.29 ± 2.80** |
| Enolase Superfamily<br>Negative Isopotential | Accuracy | 52.87 ± 35.56 | **98.50 ± 3.38** |
| | F1 | 60.75 ± 34.95 | **99.17 ± 1.88** |

Overall, DeepVASP-E substantially outperformed PCA on every dataset, indicating that cavity fields from different subfamilies could not generally be distinguished by differently located electrostatic isopotentials. Compared to PCA, another benefit of the proposed DeepVASP-E model is that it directly handles the 3D volumetric data, thus avoiding vectorization of input tensor data, which leads to certain loss of structure information.

It is also noticed that PCA’s accuracy was affected by the imbalanced size of protein subfamilies. When one protein from the chymotrypsin or muconate lactonizing enzyme was left out, the resulting classifications were near random and also affected by differences in the size of the other two subfamilies. This effect could also be seen in CNN classification, but it had a less negative effect on classification accuracy.

### Mechanism Prediction

We hypothesize that the most salient voxels identified by DeepVASP-E will be regions of electrostatic isopotential that are mechanistically involved in the specificity of the protein. This hypothesis is supported in part by the fact that DeepVASP-E can accurately classify the cavity fields from novel proteins, as demonstrated above, but we also inspected and evaluated the mechanistic accuracy of the most salient voxels.

Overall, we observed that the most salient voxels appear nearby charged amino acids that are known the affect specificity. The clearest example of this observation is the case of aspartate 189 in atlantic salmon trypsin. This amino acid has been experimentally established to play a pivotal role in the selection of positively charged substrates through electrostatic complementarity.^24^ Looking at the geometry of the negative isopotential, the most salient region for classification is a region deep in the cavity nearby aspartate 189 (Fig. 3a). It is clear that the most salient voxels in the negative isopotential are identifying a region that enables specificity in trypsin. As we look at additional voxels with diminishing salience, we can see that they emanate away from this influential region (Fig. 3b). It is clear that this region plays a crucial role in distinguishing the subfamilies of the serine proteases, and that saliency mapping is able to detect electrostatic influences on specificity.

**Fig. 3.**
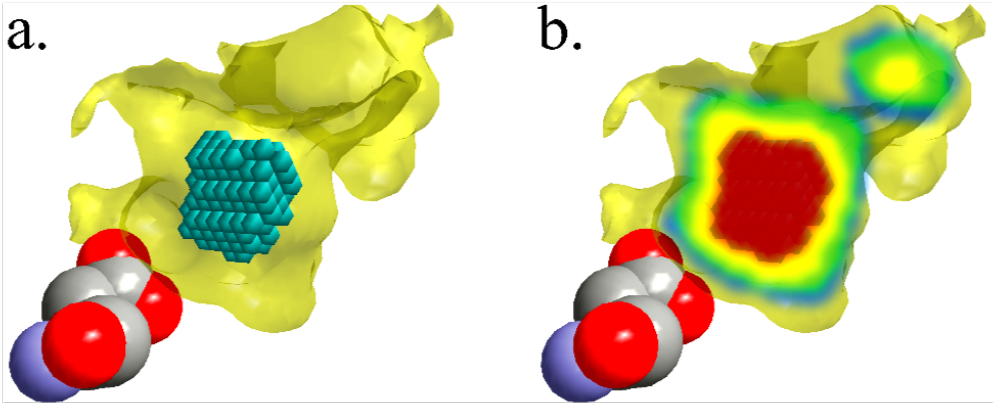
Salient Negatively Charged Regions of the Trypsin S1 Cavity. The S1 cavity of atlantic salmon trypsin (1a0j) is predominantly occupied with a negative isopotential (transparent yellow surface) generated by aspartate 189 (spheres). The most salient 150 voxels identified by DeepVASP-E are illustrated as teal cubes in **a**. This region is the area closest to D189 that maintains solvent accessibility in most conformations of the MD simulation. The next most salient groups of voxels, shown in groups of 150, are shown in **b**, in yellow, green, and blue, in descending significance.

Small regions of positive charge appear to distinguish the other subfamilies of the serine proteases. While these isopotentials are not large, they appear to support the separation of elastase from chymotrypsin. Small salient regions were often observed around the nitrogen atoms of the V216, G193 and N192 that are near the binding site. Valine 216 is known to have a steric role in elastases for excluding larger hydrophobic substrates, but not an electrostatic one.^25^ These findings illustrate that small regions of electrostatic isopotential are not random, and that they may distinguish between proteins in different subfamilies without themselves having an electrostatic role in specificity.

Among the enolase subfamily of the enolase superfamily, negatively charged isopotentials produced many salient voxels nearby Glu287, D241, and D314, which are known for stabilizing the magnesium ions necessary for dehydrating the enolase ligand. Similar salient voxels were observed near corresponding amino acids of the other members of the enolase subfamily. Among the mandelate racemaces, negatively charged isopotentials produced salient voxels nearby E247 and E317, which have a role as a general acid catalyst.^44^

Positively charged isopotentials in enolase subfamily were associated with regions of salient voxels nearby K339 and K390 in pdb 1IYX. The same amino acids in the other members of the enolase subfamily, with slightly different indices, also produced regions of salient voxels. Altogether, K339 and K390 are believed to electrostatically stabilize the carboxylate moiety of the enolase substrate.^45^ Among the mandelate racemases, salient voxels were associated with H297, K166, and K164 in pdb 1MDR. H297 and K166 are associated with proton exchange as a result of their net charge, and K164 is believed to electrostatically stabilize the carboxylate oxygen of the substrate.^44^

## 4. Conclusions

We have presented DeepVASP-E, a deep learning algorithm for detection salient features in the classification of electrostatic isopotentials within ligand binding cavities. DeepVASP-E is the first algorithm to contribute to an Analytic Ensemble strategy, by which intentionally narrow algorithms designed to predict influences on specificity can associate a biochemical mechanism with their prediction because they consider only that mechanism. DeepVASP-E focuses on electrostatic isopotentials in this manner. It can also be used to classify binding sites according to their binding preferences.

DeepVASP-E uses saliency mapping to predict regions of electrostatic potential that are important for accurate classification. We hypothesized that these regions would also be functionally significant for achieving binding specificity. In a detailed study of the amino acids nearby salient regions, we found that many of these amino acids are associated with an electrostatic role in binding specificity. We also observed that small salient regions may identify regions that assist in classification but do not contribute an electrostatic role in specificity. These findings generally support the approach of using salient regions to identify electrostatic mechanisms that influence specificity, especially if smaller salient regions could be filtered out.

Altogether, these capabilities point to applications in anticipating mutations that alter binding preferences. These include forecasting mutations that may arise in viral evolution, leading to vaccine resistance, or in protein redesign, where mutations can be suggested for altering binding specificity.

## Acknowledgements

The authors are grateful to Dr. Edward Kim for his generous advice on interpretable machine learning methods and to Mr. Desai Xie for his early work on the project. This work was funded in part by NIH Grant R01GM123131 to Brian Y. Chen. Preprint of an article submitted for consideration in Pacific Symposium on Biocomputing © [2021] World Scientific Publishing Co., Singapore, http://psb.stanford.edu/

